# An engineered glioblastoma model yields novel macrophage-secreted drivers of invasion

**DOI:** 10.1101/2023.11.18.567683

**Authors:** Erin A. Akins, Dana Wilkins, Manish K. Aghi, Sanjay Kumar

**Affiliations:** University of California, Berkeley – University of California, San Francisco Graduate Program in Bioengineering, Berkeley, CA, USA; Department of Bioengineering, University of California, Berkeley, Berkeley, CA 94720, USA; Department of Neurosurgery; University of California San Francisco (UCSF); Department of Chemical and Biomolecular Engineering, University of California, Berkeley, Berkeley, CA 94720, USA

## Abstract

Glioblastomas (GBMs) are highly invasive brain tumors replete with brain- and blood-derived macrophages, collectively known as tumor-associated macrophages (TAMs). Targeting TAMs has been proposed as a therapeutic strategy but has thus far yielded limited clinical success in slowing GBM progression, due in part to an incomplete understanding of TAM function in GBM. Here, by using an engineered hyaluronic acid-based 3D invasion platform, patient-derived GBM cells, and multi-omics analysis of GBM tumor microenvironments, we show that M2-polarized macrophages stimulate GBM stem cell (GSC) mesenchymal transition and invasion. We identify TAM-derived transforming growth factor beta induced (TGFβI/BIGH3) as a pro-tumorigenic factor in the GBM microenvironment. In GBM patients, BIGH3 mRNA expression correlates with poor patient prognosis and is highest in the most aggressive GBM molecular subtype. Inhibiting TAM-derived BIGH3 signaling with a blocking antibody or small molecule inhibitor suppresses GSC invasion. Our work highlights the utility of 3D *in vitro* tumor microenvironment platforms to investigate TAM-cancer cell crosstalk and offers new insights into TAM function to guide novel TAM-targeting therapies.

## MAIN

Glioblastoma (GBM) is the most common, lethal, and aggressive form of primary brain cancer with a medium survival rate of approximately 15 months.^1^ The highly invasive nature of GBM complicates surgical resection and promotes recurrence, yet it has been challenging to target invasion therapeutically due to an incomplete understanding of the underlying cellular and molecular mechanisms. Cell invasion is profoundly influenced by interactions between cancer cells and the tumor microenvironment (TME), which consists of non-neoplastic cells as well as extracellular matrix (ECM).^2^ Immune cells have emerged as pivotal TME players in tumor development, making them highly attractive targets for therapeutic intervention.^3,4^ GBMs exhibit a unique immune landscape dominated by tumor-associated macrophages (TAMs), which are composed of multiple subpopulations, including bone marrow-derived (BMD) macrophages and brain-resident microglia.^5,6^ TAMs can comprise up to 40% of the total cells in gliomas and their accumulation is highest in patients with the most aggressive GBM subtype.^6–9^ TAMs are proposed to actively regulate multiple cellular properties relevant to tumor progression including stemness, proliferation, survival and migration, but the clinical impact of TAM-tumor interactions appears to be strongly influenced by characteristics of the tumor (location, stage, subtype) as well as macrophage cell phenotypes. ^10–15^

Macrophage phenotype and function is defined by a polarization state, which has traditionally been described using the M1/M2 polarization model.^16,17^ TAMs are often associated with the anti-inflammatory M2 polarization state due to their high expression of anti-inflammatory cytokines, scavenger receptors, pro-angiogenetic factors and proteins involved in ECM organization and remodeling.^18^ The pro-inflammatory M1-like state is more commonly associated with anti-tumor functions such as antigen-presentation and co-stimulation. Although the M1/M2 classification system has proven to be valuable for in vitro studies, TAM polarization state is increasingly viewed as much more nuanced than the binary M1/M2 classification would suggest, and human GBM TAMs have been found to display both M1 and M2 gene signatures.^19–22^

An important and ongoing barrier to understanding TAM regulation of GBM invasion is the lack of in vitro platforms that capture key geometric and mechanical aspects of the GBM TME. Advances in biomaterials and tissue engineering have facilitated the development of 3D TME models.^23,24^ Hydrogel-based systems are frequently used as matrices for 3D cell culture, and among these systems, hyaluronic acid (HA) stands out as a valuable biomaterial specifically for modeling the brain TME.^25^ We and others have used HA-based hydrogel platforms to identify and mechanistically dissect key pathways driving GBM invasion, metabolism, and therapeutic resistance.^26–32^ Despite the rapid expansion in 3D brain TME models, integration of macrophages and other immune cells in these platforms remains limited. ^33,34^

Here we utilize a 3D HA-based hydrogel platform to test the hypothesis that TAMs establish a pro-invasive microenvironment that facilitates GBM invasion. We first investigate the influence of macrophage polarization state on GBM spheroid invasion across a panel of patient-derived GBM cells. We find that factors secreted by M2-polarized macrophages induce GBM invasion. Using a microscale invasion platform that enables regional dissection and characterization of invasive cells, we identify macrophage-induced transcriptional changes in invasive GBM cells. We then characterize the macrophage secretome across polarization states using mass spectrometry, perform a GBM-TAM interaction analysis to identify receptor-ligand pairs driving invasion, and prioritize putative pro-invasive TAM-secreted ligands using single-cell RNA sequencing (scRNA-seq) datasets of human GBM TAMs. We identify two novel TAM-derived secreted factors, BIGH3 and S100A9, that stimulate GSC invasion. We demonstrate that targeting BIGH3 and downstream mammalian target of rapamycin (mTOR) signaling reduces invasion. Overall, our study highlights the utility of engineered 3D platforms to investigate TAM-cancer cell interactions and uncovers novel molecular targets.

## Results

### Establishment of a brain-inspired hydrogel platform to study TAM-GBM interactions

We engineered a 3D in vitro brain tumor model using our previously published HA-RGD hydrogel formation.^35^ (**Fig. 1a; Ext. Fig. 1a**) To study the influence of macrophages on GBM invasion, we performed co-culture tumorsphere invasion assays by encapsulating patient-derived GBM stem cells (GSCs) in HA-RGD hydrogels with THP-1-derived M2-polarized macrophages distributed throughout the hydrogel (**Fig. 1b-c; Ext. Fig. 1b-e**). As monocultures, GSC-295 spheroids are not invasive; however, adding M2-polarized macrophages resulted in a pro-invasive GSC phenotype characterized by long, thin protrusions in the matrix, which we quantified by calculating invasive cell area (*See Methods*). To test whether M2 macrophages influence invasion through direct or paracrine mechanisms, we performed indirect co-culture assays using M2-polarized macrophage conditioned media (M2 CM). Treatment with M2 CM recapitulated the invasive phenotype seen in direct co-culture, suggesting that GSC invasion did not require macrophage-GSC physical contact (**Fig. 1d,e)**.

**Figure 1:**
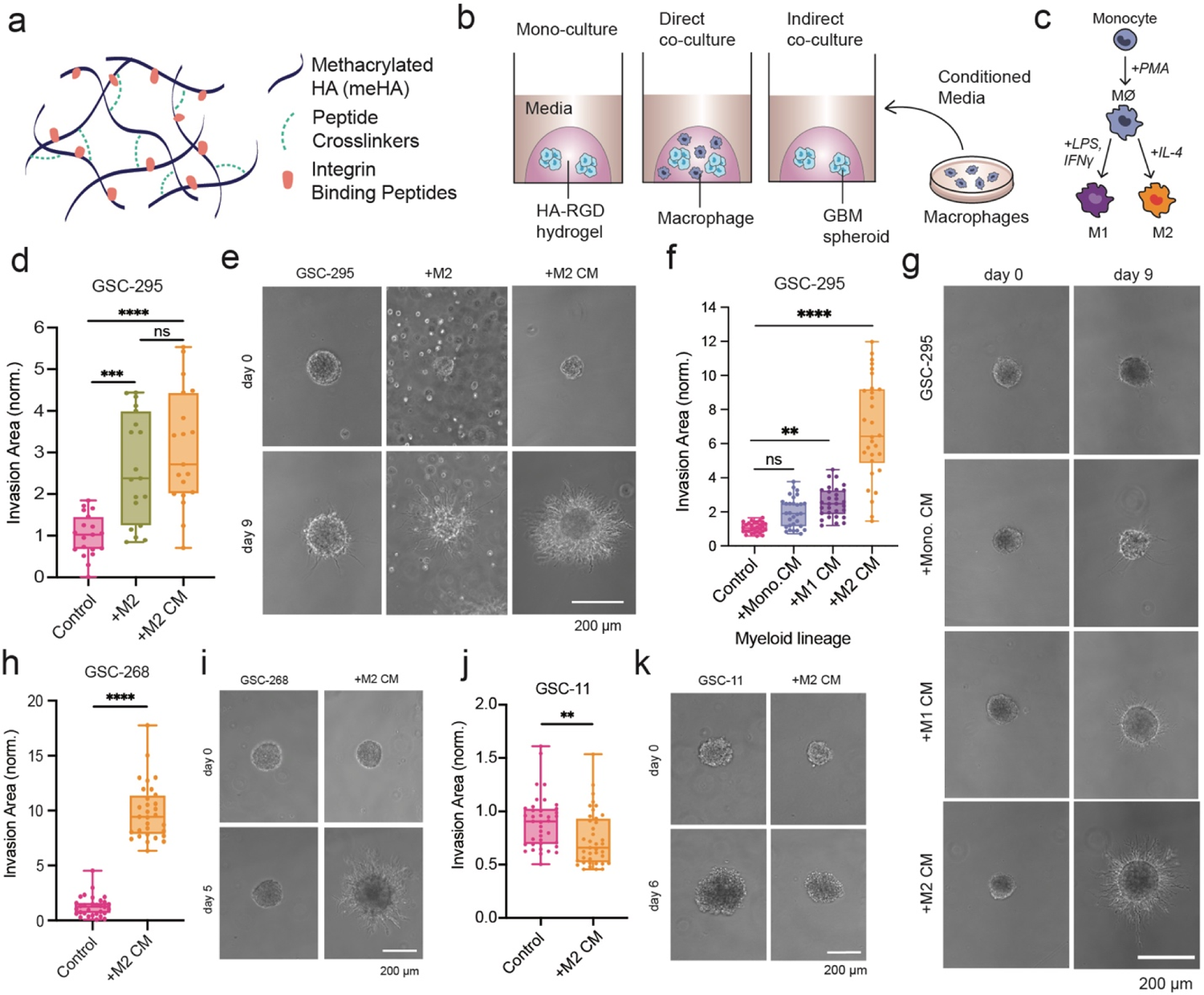
An engineered biomaterials platform to enable investigation of GBM-macrophage crosstalk. **a**, Schematic of HA-based hydrogel. **b,** Schematic of mono- or co-culture platforms to study GBM invasion. **c,** Schematic of THP-1-derived monocyte to macrophage differentiation and M1/M2 polarization protocol. **d,e,** GSC-295 invasion assay with direct M2 macrophage co-culture and M2 macrophage conditioned media (CM) (**d**) quantification and (**e**) representative phase images. **f,g,** GSC-295 invasion assay with CM from monocytes (mono.), M1 macrophages, and M2 macrophages (**f**) quantification and (**g**) representative phase images. **h,i,** GSC-268 invasion assay with M2 CM (**h**) quantification and (**i**) representative phase images. **j,k,** GSC-11 invasion assay with M2 CM (**j**) quantification and (**k**) representative phase images.

To examine the effect of polarization state on macrophage-GSC interactions, we treated GSC-295 spheroids with CM collected from monocytes and M1-polarized macrophages. GSC-295 spheroids cultured in monocyte CM (mono. CM) and M1 macrophage CM (M1 CM) were less invasive than GSC-295 spheroids cultured in M2 CM (**Fig. 1f,g**). These results suggest that in our system, M2 macrophages are the predominant source of pro-invasive factors.

### GBM-Macrophage interaction varies across a panel of patient-derived GBMs

To determine whether M2 CM stimulated invasion of additional patient-derived GSCs, we performed tumorsphere invasion assays using a previously characterized panel of patient-derived GSCs spanning three subtype classifications (mesenchymal, proneural, classical).^36^ The influence of M2 CM on spheroids appeared to be cell line-specific, but the majority of GSC lines (5/6) were highly invasive when treated with M2 CM (**Fig. 1h,i; Ext. Fig. 2a-f).** Interestingly, GSC-11 spheroids decreased in area when treated with M2 CM (**Fig. 1j,k).** We confirmed the polarization-specific response and found that like

**Figure 2:**
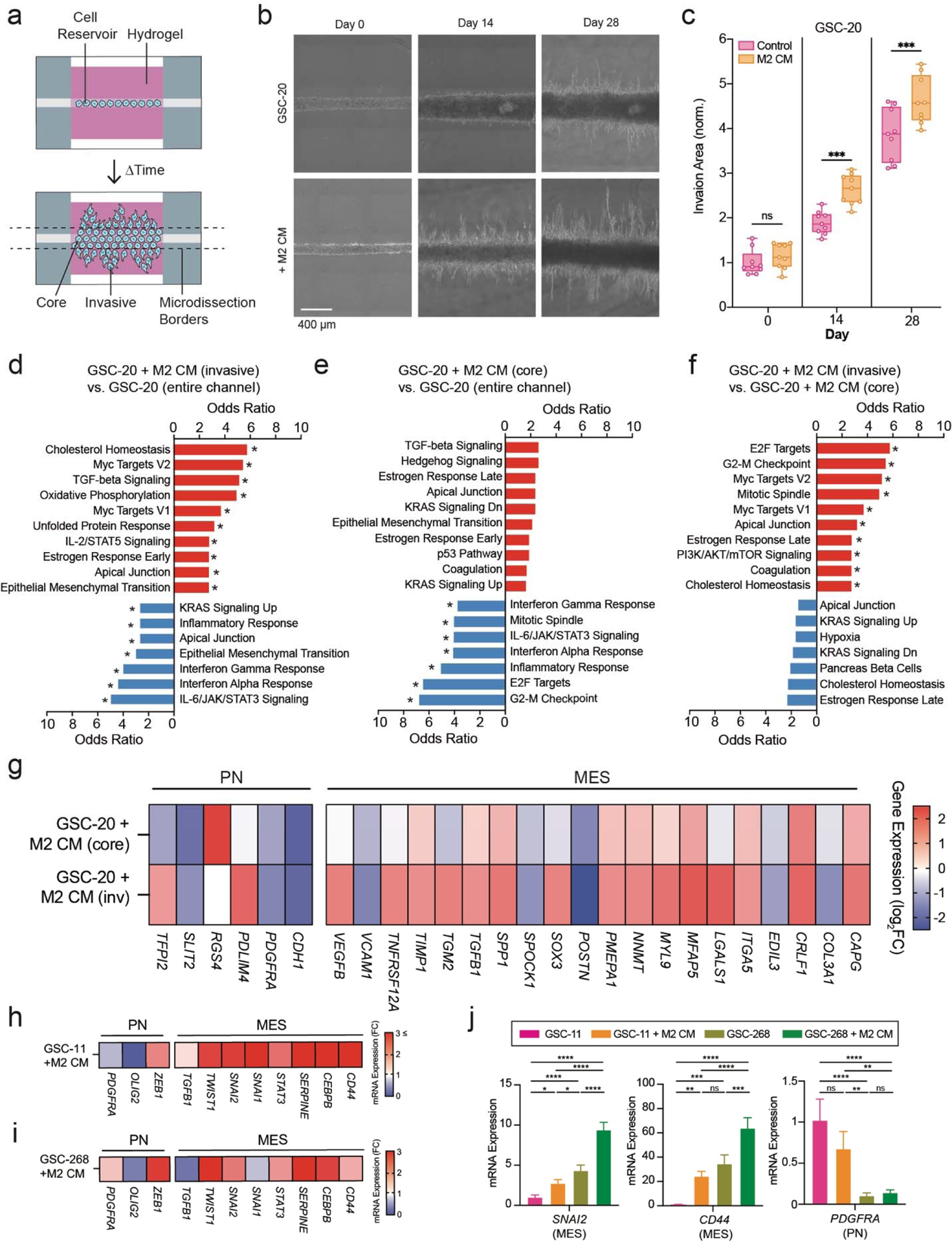
GSC transcriptional changes induced by M2 CM-mediated invasion. **a**, Schematic of HA-based invasion device and microdissection borders (dashed lines). **b,c,** GSC-20 invasion devices with M2 CM (**b**) representative phase images and (**c**) quantification. **d-f,** Cellular pathway (MSigDB^1^) enrichment analysis of differentially expressed genes. Pathways, odds ratio and statistical significance (\**P*_adj_ < 0.05) calculated from Enrichr^2–4^ between (**d**) GSC-20 invasive fraction of M2 CM devices and entire channel of GSC-20 control devices, (**e**) GSC-20 core fraction of M2 CM devices and entire channel of GSC-20 control devices and (**f**) GSC-20 invasive fraction of M2 CM devices and GSC-20 core fraction of M2 CM devices. **g,** Heat map showing GSC-20 relative gene expression levels of mesenchymal (MES) and proneural (PN) markers obtained by bulk RNA-seq of cells isolated from invasion devices. Expression levels normalized to GSC-20 devices in control media (entire channel). **h,i,** Heat maps showing relative gene expression levels of MES and PN markers obtained by qPCR of cells isolated from spheroid invasion assays. Expression levels are normalized to GSCs in control media (**h**) GSC-11 and (**i**) GSC-268. **j,** Relative gene expression levels of PDGFRA, SNAI2, and CD44 across GSC lines and culture conditions. Expression levels for each gene are normalized to GSC-11 in control media.

GSC-295, M2 CM drives GSC-268 invasion greater than mono. CM or M1 CM (**Ext. Fig. 2g,h**).

To test whether M2 CM stimulates invasion in patient-derived cells that have not been enriched for GSCs, we performed tumorsphere invasion assays with two GBM patient-derived xenograft lines (GBM-43 and GBM-6).^37^ Interestingly, GBM-43 spheroids decreased invasion when cultured with M2-polarized macrophages or in M2 CM (**Ext. Fig. 2i,j**). A similar trend was seen in GBM-6 spheroids, although the difference was not statistically different (**Ext. Fig. 2k,l**).

Using GSC-268 spheroids and varied concentrations of M2 CM, we found that M2 CM stimulates invasion in a dose-dependent fashion (**Ext. Fig. 2m**). To test whether the increased invasion was connected to differences in proliferation, we quantified the percentage of KI-67 positive cells following an invasion assay and found no difference in the percentage of KI-67 cells between GSCs cultured in control GSC media and M2 CM (**Ext. Fig. 2n,o**).

### M2-polarized macrophage secreted factors shift GSC transcriptional profiles towards invasion- and mesenchymal-associated signatures

To investigate how M2 CM alters the transcriptional profile of GSCs, we performed bulk RNA-sequencing (RNA-seq) on GSCs in hydrogels cultured in GSC maintenance (control) media or M2 CM. We utilized our previously developed microchannel invasion platform, which consists of a cylindrical cell reservoir channel embedded within a 3D hydrogel and enables the physical separation of highly invasive and non-invasive (core) cell fractions from a single device (**Fig. 2a**).^28,32^ As expected, GSC-20s in M2 CM-treated devices were more invasive than GSC-20s in control devices (**Fig. 2b-c**).

Following the invasion assay, we micro-dissected the M2 CM-treated devices to isolate the highly invasive GSCs. We performed differential gene expression analysis between cells isolated from control devices, invasive cells isolated from M2 CM-treated devices, and core cells isolated from M2 CM-treated devices. We found invasive and core populations separated along the first principal component (PC1), indicating maximum variance along the invasive phenotype. The second principal component (PC2) generally separated GSCs based on media conditions (M2 CM vs. control media) (**Ext. Fig. 3a**). We identified ∼1000 differentially expressed genes (DEGs) between each sample comparison illustrated in the heat map and volcano plots (**Ext. Fig. 3b-f, Supp Table 1**).

**Figure 3:**
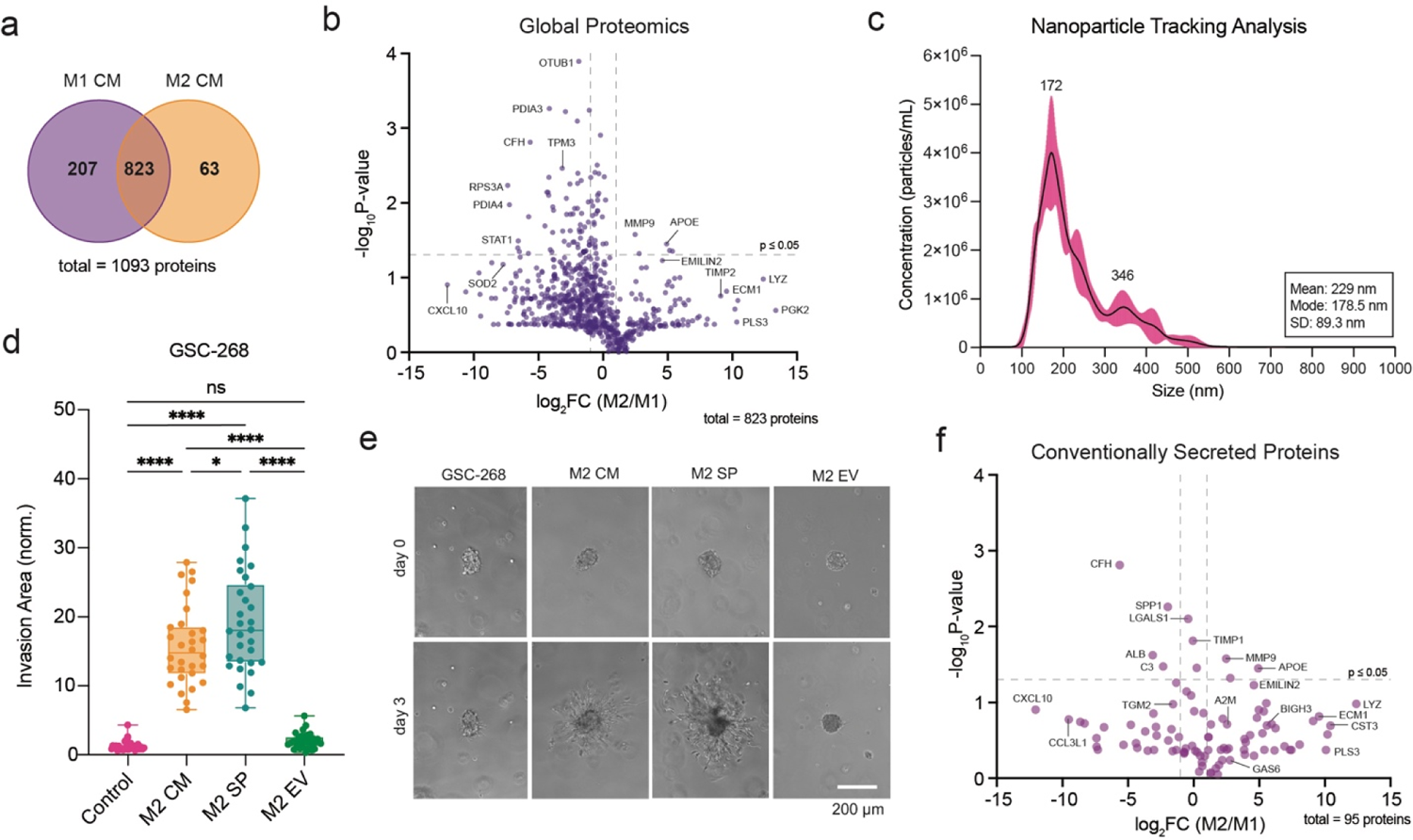
Mass Spectrometry-based identification of macrophage secreted proteins. **a**, Venn Diagram illustrating proteins identified by mass spectrometry in M1 and M2 CM. **b,** Volcano plot of proteins identified by mass spectrometry in both M1 and M2 CM. **c,** Nanoparticle tracking analysis (NTA) of extracellular vesicles (EV) isolated by ultracentrifugation from M2 CM. **d,e,** GSC-268 invasion assay with M2 CM, M2 soluble proteins (SP) and M2 EV (**d**) quantification and (**e**) representative phase images. **f,** Volcano plot of conventionally secreted proteins identified by mass spectrometry in both M1 and M2 CM.

To identify pathways associated with the invasive phenotypes in response to M2 CM, DEGs were used as inputs for pathway enrichment analysis (**Fig. 2d-f**). M2 CM-treatment led to downregulation of pathways related to Interferon Gamma Response, Interferon Alpha Response, Inflammatory Response and IL-6/JAK/STAT3 Signaling, and upregulation of mesenchymal (MES) pathways including TGF-β Signaling, Apical Junction and Epithelial Mesenchymal Transition (EMT) (**Fig. 2d,e**). Invasive GSCs were associated with enrichment pathways related to Cholesterol Homeostasis, MYC Targets V1/V2, and Apical Junction (**Fig. 2d,f**).

Interestingly, MES pathways (TGF-β Signaling, Apical Junction and EMT) were upregulated in both the invasive and core populations of M2 CM-treated devices, suggesting that M2 CM treatment induces global upregulation of MES-related genes in GSCs. We further examined bulk RNA-seq expression levels of MES-associated genes across device fractions and culture conditions (**Fig. 2g**). We also included DEGs associated with the less aggressive proneural (PN) subtype, which has better overall patient survival compared to the MES subtype.^8^ Compared to GSCs isolated from control devices, GSCs cultured in M2 CM had elevated expression of MES genes and downregulated expression of PN genes. The gene expression changes were seen in both M2 CM-treated core and invasive cells but were larger in the M2 CM-treated invasive cell population. Treatment with M2 CM also led to differential expression of cadherins, including the characteristic MES-associated downregulation of E-cadherin (*CDH1*) (**Ext. Fig. 3g**).^38^

We next explored whether M2 CM induced upregulation of MES genes across multiple patient-derived GSCs. GSC-11 spheroids, which do not invade when cultured in M2 CM, displayed an upregulation of MES-associated genes (*CD44, CEBPB, SERPINE, SNAI1*) and a downregulation of PN-associated genes (*OLIG2, PDGFRA*) relative to GSC-11 spheroids in control media (**Fig. 2h**). GSC-268 spheroids, which do invade when cultured in M2 CM, also displayed an upregulation of MES-associated genes (**Fig. 2i**). To better understand the association of MES and PN gene signatures with invasion, we compared gene expression levels for *PDGFRA* (PN marker), *SNAI2* (MES marker) and *CD44* (MES marker) between GSC-11 and GSC-268. The relative expression of MES marker genes is generally lower in GSC-11 spheroids than GSC-268 spheroids regardless of media condition (**Fig. 2j)**. Additionally, GSC-11 spheroids exhibited higher gene expression levels of the PN marker *PDGFRA* relative to GSC-268 spheroids, independent of media condition (**Fig. 2j**).

### Identification of macrophage-derived pro-invasive proteins using mass spectrometry

To directly identify soluble factors secreted by TAMs that may induce MES genes and drive GSC invasion, we characterized macrophage CM with global proteomics. Using size-based exclusion filtration and heat treatment of M2 CM, we first determined that the pro-invasive factor in M2 CM was greater than 10 kDa (**Ext. Fig. 4a**) and heat-sensitive (**Ext. Fig. 4b**). From these data, we hypothesized that the pro-invasive factor was a protein and performed mass spectrometry to identify secreted proteins that were more abundant in M2 CM than M1 CM. Mass spectral analysis identified 1093 total proteins, with 207 proteins detected only in M1 CM, 63 proteins detected only in M2 CM, and 823 proteins detected in both M1 and M2 CM (**Fig. 3a,b; Ext. Fig. 4d; Dataset 2**).

**Figure 4:**
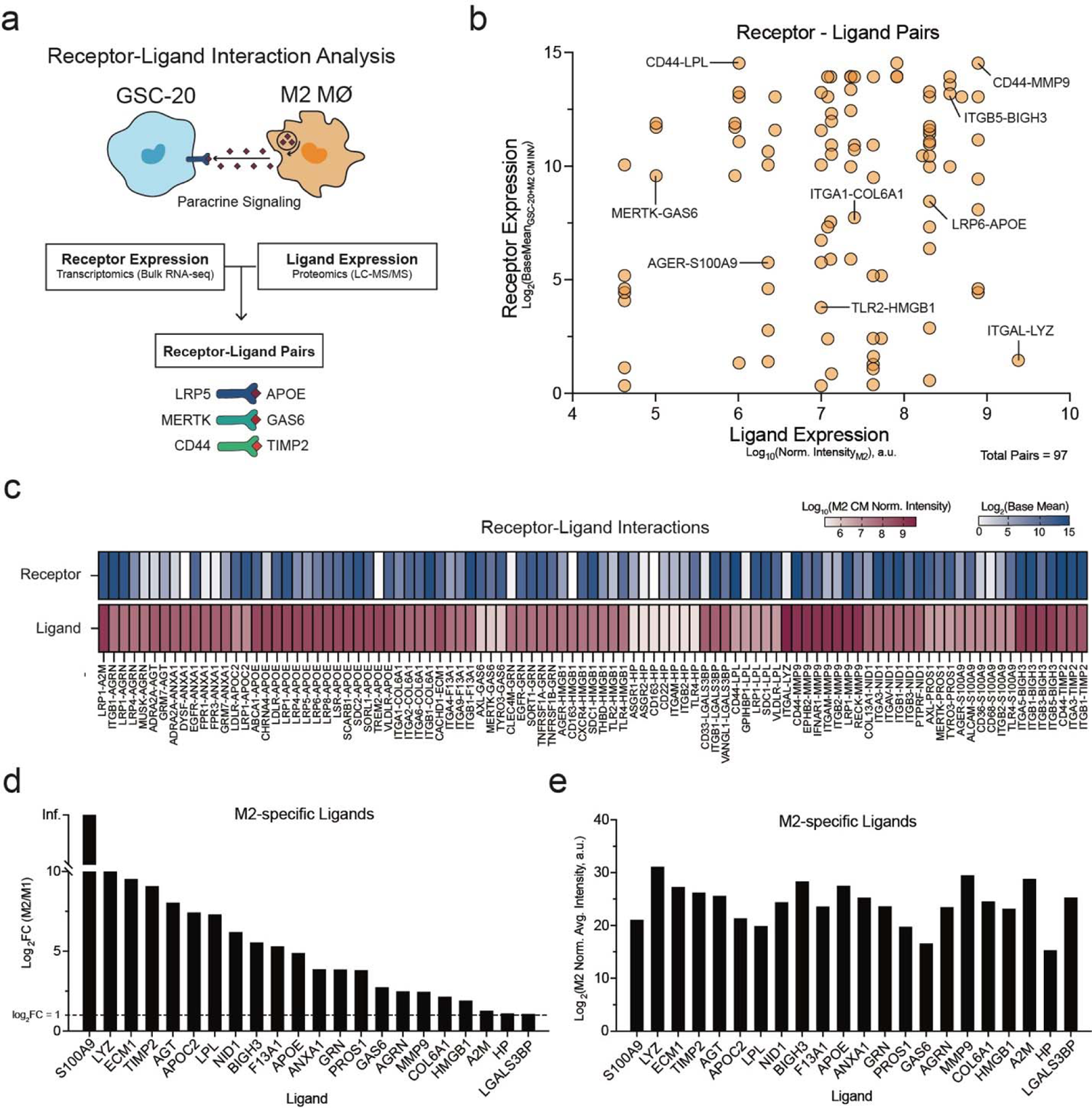
Paracrine receptor-ligand interaction analysis of GSCs and M2 macrophages. **a**, Schematic of Receptor-ligand interaction analysis workflow. **b,** Quantification of receptor-ligand pair according to ligand expression in M2 CM (x-axis) and receptor expression in GSCs (y-axis). **c,** Heat map of receptor expression in GSCs and ligand expression in M2 CM for each receptor-ligand pair. **d,e,** Bar graphs of ligands detected in M2 CM emerging from receptor-ligand analysis graphed to illustrate (**d**) average protein abundance in M2 CM compared to M1 CM and (**e**) average relative protein abundance obtained from mass spectrometry.

As expected, M1 CM had higher abundance of pro-inflammatory proteins (CXCL8, CCL3, CXCL10, CCL5, IFI30) while M2 CM had higher abundance of anti-inflammatory and ECM remodeling proteins (TIMP2, BIGH3, ECM1, ADAM10, CSTHB) (**Ext. Fig. 4e**). Beyond these conventionally secreted proteins, mass spectrometric analysis identified various unconventionally secreted proteins such as OTUB1, PDIA3, and PGK2 (**Fig. 3b**). We hypothesized that these detected proteins could have been secreted through extracellular vesicles (EVs). We isolated and purified EVs from M2 CM and performed a tumorsphere invasion assay with full M2 CM, M2 EVs and EV-free M2 CM, referred to as M2 soluble protein fraction (M2 SP). GSC-268 spheroids invaded when treated with M2 SP and M2 CM, but were not invasive when treated with M2 EVs (**Fig. 3c-e**). GSC-295 spheroids displayed a similar trend and were only invasive when treated with M2 CM or M2 SP (**Ext. Fig. 4f,g**). From these results, we hypothesized that the pro-invasive factor was not EV-bound and computationally filtered our list of identified proteins to include only conventionally secreted proteins (**Fig. 3f**).^39^

### GSC-macrophage interaction analysis identifies putative receptor-ligand pairs driving invasion

To identify specific receptor-ligand pairs that may be driving invasion, we created a resource of inferred paracrine crosstalk by mapping the expression of GSC receptors to that of their cognate ligands detected in macrophage CM (**Fig. 4a**). We identified 97 receptor-ligand pairs of which the receptor is expressed by GSC-20s and the ligand is present in M2 CM (**Fig. 4b; Dataset 3**). The 97 pairs included 65 unique receptors and 22 unique ligands (**Fig. 4c**) and although all ligands were detected at higher abundance in M2 CM than M1 CM (**Fig. 4d**), their relative expression in M2 CM varied (**Fig. 4e**).

### Candidate M2 macrophage secreted factors are abundant in human GBM TME and preferentially expressed by TAMs

We next investigated the relevance of candidate ligands to human GBM tumors by re-analyzing a publicly available single cell RNA-seq (scRNA-seq) dataset (ref. ^40^). Since all but one of the 22 ligands identified from our receptor-ligand interaction analysis was a DEG, we focused our analysis on ligands that were predominantly expressed by TAM clusters: *ECM1, A2M, APOE, GAS6, BIGH3, S100A9, MMP9, GRN, APOC2, LYZ* (**Fig. 5a**). To predict whether any of these ligands were specific to microglia or BMD TAMs we probed a subclustered version of the same dataset. Every ligand was identified in at least 1 TAM cluster but some ligands were preferentially expressed by either microglia or BMD TAMs (**Fig. 5b**). Since the majority of TAMs in GBM are of monocytic origin, and CM from polarized human microglia did not drive GSC invasion (**Ext. Fig. 5a,b**), we prioritized ligands predominantly expressed by BMD TAMs (*BIGH3*, *S100A9* and *LYZ*). We validated these findings using a second publicly available human GBM scRNA-seq dataset (ref. ^41^) and found that the top three DEGs upregulated in BMD TAMs were again *BIGH3, S100A9 and LYZ* (**Fig. 5c**).

**Figure 5:**
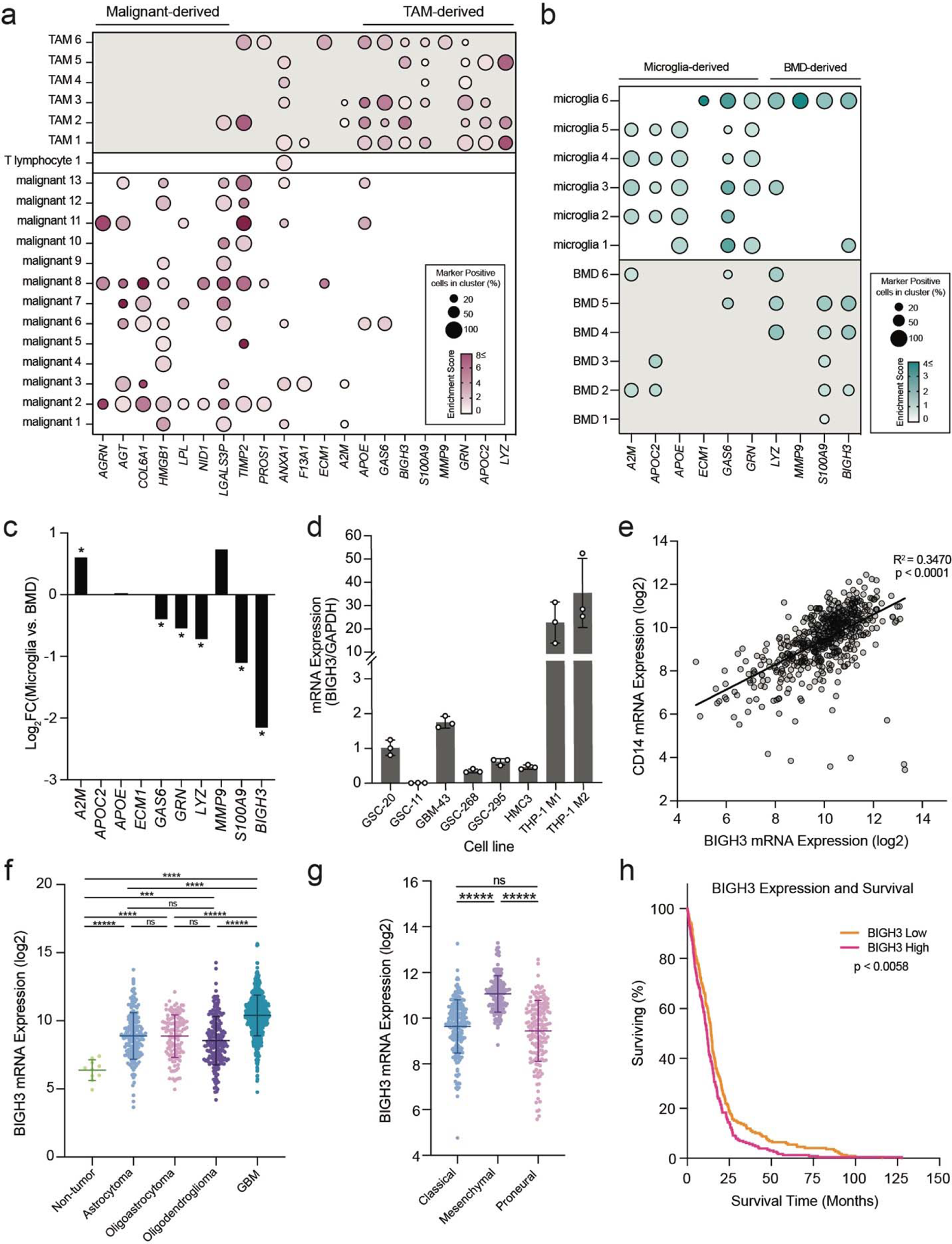
Integration of M2 macrophage-secreted ligands with published transcriptomic datasets. **a,b**, Dot plot depicting gene expression profiles of M2 macrophage-secreted ligands identified by receptor-ligand analysis. Human GBM scRNA-seq dataset was obtained from Cui, et al.^5^ Plots include (**a**) gene expression of all predicted M2 macrophage ligands across all GBM tumor cell type clusters and (**b**) gene expression of top 10 predicted M2 macrophage ligands across only GBM TAM clusters. **c,** Differential gene expression analysis between human GBM bone-marrow-derived TAMs and microglia TAMs. Genes with log_2_FC > 1 are expressed higher in microglia TAMs. Human scRNA-seq dataset was obtained from Müller, et al.^6^ **d,** Relative BIGH3 mRNA expression obtained from qPCR across a panel of cell lines. **e-h**, Gene expression data from TCGA database queried and downloaded using GlioVis^7^ data portal for visualization and analysis of brain tumor expression datasets. (**e**) Simple Linear Regression plot and analysis of CD14 and BIGH3 gene expression in GBM tumors. BIGH3 mRNA expression (log2) across (**f**) glioma tumor stage and (**g**) GBM molecular subtype. (**h**) Kaplan–Meier survival correlating with high or low levels of BIGH3 mRNA with median expression level used as cutoff. *P* values determined by log-rank Mantel–Cox comparison (n=262 patients in the high expression group, and n=263 patients in the low expression group).

We focused further mechanistic investigation on transforming growth factor beta induced (TGFBI/BIGH3), as this secreted extracellular protein was one of the top ligands emerging from the scRNA-seq analysis. We measured BIGH3 mRNA expression levels across a panel of human cells and found that M2-polarized macrophages had the highest BIGH3 mRNA expression levels compared to GBM and microglia lines (**Fig. 5d**).

To determine if BIGH3 mRNA expression varied by glioma grade, GBM subtype, and GBM patient survival, we used GlioVis data portal for visualization and analysis of brain tumor expression datasets.^42^ In patients with GBM, we found a positive linear correlation between mRNA expression levels of BIGH3 and the monocytic marker CD14 (**Fig. 5e**). BIGH3 mRNA expression level was highest in GBM tumors compared to lower grade gliomas and non-tumor regions (**Fig. 5f**) and varied by GBM molecular subtype, with highest expression in patients classified as MES subtype. Furthermore, high BIGH3 gene expression correlated with shorter patient survival (**Fig. 5h**).

### TAM-secreted BIGH3 promotes GSC invasion

We next tested whether recombinant human BIGH3 (rhBIGH3) was sufficient to drive GSC invasion. We found that while rhBIGH3 induced GSC-268 spheroid invasion (**Fig. 6a-b**), rhBIGH3 interestingly had no effect on GSC-11 spheroids, which also did not invade with M2 CM (**Fig. 6c-d**). To test the functional contributions of BIGH3 in M2 CM-mediated invasion, we performed invasion assays using a blocking antibody targeting BIGH3 (α-BIGH3). GSC-268 spheroids cultured in M2 CM and α-BIGH3 were less invasive than spheroids cultured in M2 CM with IgG isotype control antibody (**Fig. 6e-f**).

**Figure 6:**
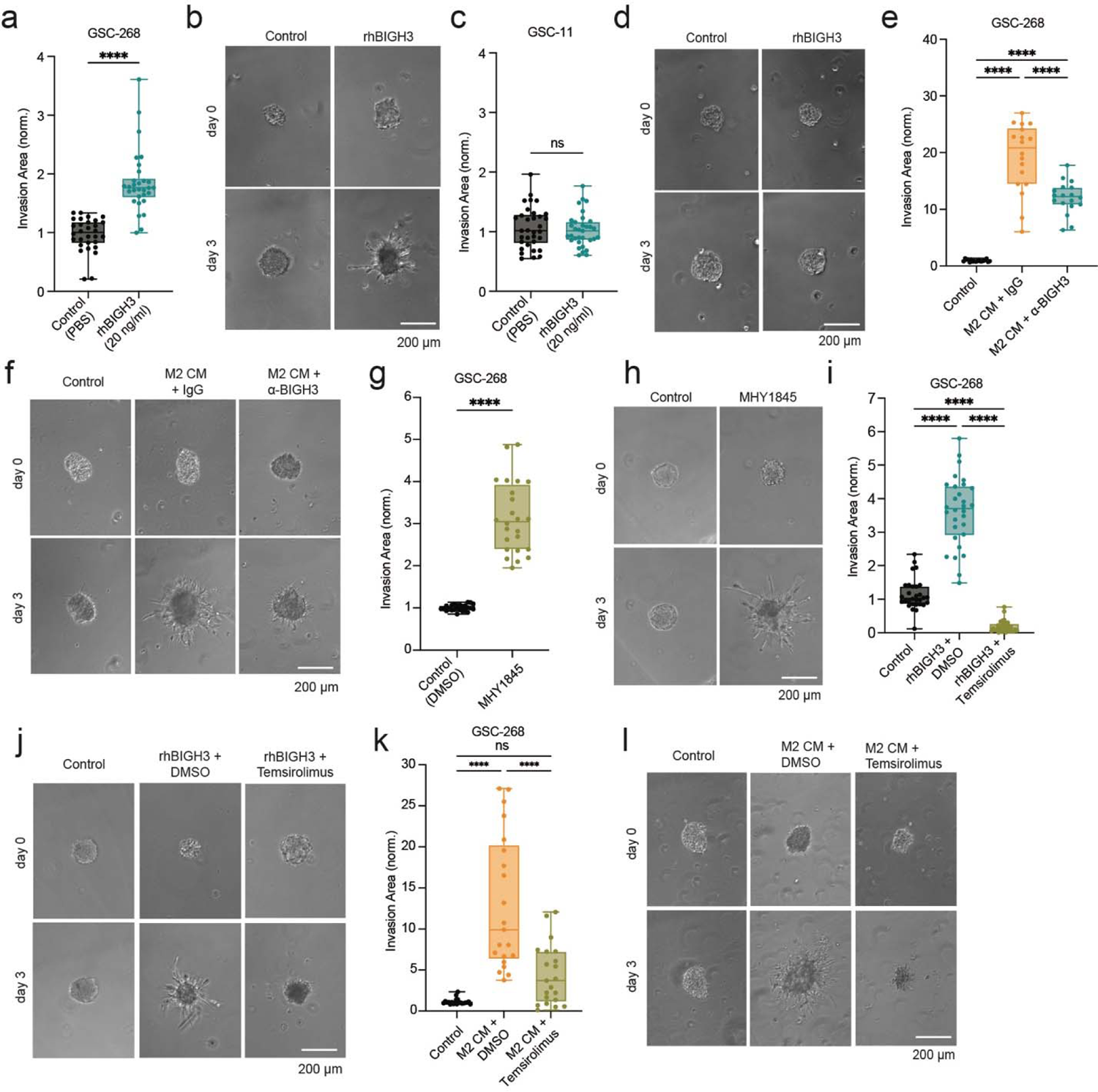
M2 macrophage-induced GSC invasion is mediated by secreted BIGH3 and is associated with active mTOR pathway. **a,b**, GSC-268 invasion assay with 20 ng/ml rhBIGH3 (**a**) quantification and (**b**) representative phase images. **c,d,** GSC-11 invasion assay with 20 ng/ml rhBIGH3 (**c**) quantification and (**d**) representative phase images. **e,f,** GSC-268 invasion assay with M2 CM neutralized with 10 µg/ml IgG (rabbit) or 10 µg/ml anti-BIGH3 (**e**) quantification and (**f**) representative phase images. **g,h,** GSC-268 invasion assay with 10 µM mTOR activator MHY1845 (**g**) quantification and (**h**) representative phase images. **i,j,** GSC-268 invasion assay with 20 ng/ml rhBIGH3 and 10 µM mTOR inhibitor Temsirolimus (**i**) quantification and (**j**) representative phase images. **k,l,** GSC-268 i nvasion assay with M2 CM and 10 µM mTOR inhibitor Temsirolimus (**k**) quantification and (**l**) representative phase images.

We next explored another top hit from our multi-omics analysis: S100 calcium binding protein A9 (S100A9). GSC-268 spheroids treated with recombinant human S100A9 (rhS100A9) exhibited increased invasion relative to spheroids cultured in control media and this interaction was inhibited using the small molecule S100A9 inhibitor, ABR-238907 (ABR) (**Ext. Fig. 5c-d**). Interestingly, ABR did not change the invasive capacity of GSC-268 spheroids cultured in M2 CM, so although S100A9 stimulates invasion in our system, it does not appear to be a targetable pro-invasive secreted factor in M2 CM (**Ext. Fig. 5e-f**).

We explored the intracellular signaling pathways activated by BIGH3 that could be driving invasion. BIGH3 is a secreted ECM protein, and its expression is regulated by transforming growth factor beta (TGF-β) signaling.^43^ According to our bulk RNA-seq analysis, TGF-β signaling and *TGF*β*1* are both upregulated in highly invasive GSCs (**Ext. Fig. 5g**). To test the possibility that BIGH3 and M2 CM promotes invasion through TGF-β signaling, we directly induced TGF-β signaling with recombinant human TGFβ1 (rhTGFβ1). The addition of rhTGFβ1 did not change the invasive capacity of GSC-268 spheroids suggesting that the induction of TGF-β signaling with rhTGFβ1 is not sufficient to drive GSC invasion (**Ext. Fig. 5h-i**).

Another pathway upregulated in highly invasive GSCs was the P13K/AKT/mTOR signaling pathway, which is abnormally activated in tumors to promote growth, metastasis and invasion.^44^ We first asked if mTOR activation was sufficient to drive invasion and found that inducing mTOR signaling with an mTOR activator (MHY1845) induced GSC-268 spheroid invasion (**Fig. 6g-h**). We next tested whether BIGH3-promoted GSC invasion was mediated by mTOR signaling. Indeed, inhibiting mTOR (Temsirolimus) slowed down rhBIGH3-mediated GSC invasion as GSC-268 spheroids treated with rhBIGH3 and Temsirolimus were significantly less invasive than spheroids treated with rhBIGH3 and vehicle control (DMSO) (**Fig. 6i-j**). We then determined what portion of M2-mediated GSC invasion was driven by mTOR by testing the ability of Temsirolimus to block M2 CM-mediated GSC invasion. We found that GSC-268 spheroids treated with M2 CM and Temsirolimus were significantly less invasive than spheroids treated with M2 CM and vehicle control (DMSO) (**Fig. 6k-l**). Inhibition of mTOR signaling through Temsirolimus blocked BIGH3- and M2 CM-mediated GSC invasion, suggesting that in our system, M2 CM and BIGH3 stimulate invasion through the mTOR pathway.

## Discussion

Tumor-immune cell interactions within the GBM TME represent a promising and largely untapped set of targets tor limiting disease progression and improving patient outcome. However, progress in identifying these targets is limited by a lack of physiologically mimetic culture platforms that support both tumor cell invasion and incorporation of TAMs. We address this need by developing 3D HA models that enable unbiased multi-omics analysis on GBM and TAM populations, which allowed us to identify pro-invasive TAM-secreted factors. We functionally validated two TAM-secreted ligands, BIGH3 and S100A9. BIGH3 has recently been recognized as a key component of the tumor ECM with both tumor suppressor and tumor promoter roles, depending on tumor type and stage.^45^ While BIGH3 has been proposed as a potential MES subtype signature gene based in part on its ability to promote growth and motility of continuous GBM cell lines, the cellular source of BIGH3 in tumors is unclear.^46,47^ Similarly, S100A9 overexpression has been documented in various human cancers and linked to TME immunosuppression and therapeutic resistance.^48–52^ Our work provides additional evidence that BIGH3 and S100A9 are active drivers of GBM invasion and suggests that they are primarily derived from the BMD TAM population.

Our findings begin to deconstruct complex TAM-GBM cell interactions and support ongoing therapeutic efforts aimed at repolarizing M2 macrophages towards M1-like polarization states, as well as offering novel therapeutic targets that could prevent the potent effects of M2 macrophages on GSCs.^53–56^ While there have been reports of high M2 TAM accumulation in MES tumors, the mechanism behind this correlation remains unclear, although it has been proposed that MES tumors attract M2 TAMs.^8,57^ Our work expands on prior studies investigating the relationship between TAM abundance and GBM subtype by proposing the possibility that M2 TAM secreted factors directly induce MES transition in tumor cells. TAMs and GSCs may potentially form a bidirectional feedback loop, where M2 TAMs induce a highly invasive MES state in GSCs, leading to further M2 polarization of TAMs. This notion is further supported by our finding that high expression of MES gene signatures correlates with highly invasive behavior, a connection previously demonstrated in breast cancer but not fully elucidated in brain tumors.^58–60^ In this study, the mTOR pathway emerged as a targetable regulator of GSC-TAM interactions and GSC invasion. Although the exact mechanisms connecting BIGH3, mTOR signaling and MES transition remain unclear, BIGH3 has several known integrin binding motifs.^61,62^ and mTOR activation has been shown to occur through integrin-dependent mechanisms.^63,64^ Given the importance of mTOR signaling in cell growth, proliferation, metabolism, and survival, this pathway is potential therapeutic target in many cancers and our data supports ongoing exploration of mTOR pathway inhibition as combination therapy for patients with GBM.^44^

By exploiting engineered 3D TME models to investigate functional contributions of the TAM secretome on GBM invasion, we show that the effects of TAM secretome on GBM are context-specific and vary by macrophage ontogeny and polarization state, as well as GBM stemness and GSC molecular subtype. BIGH3 and other TAM-secreted factors merit further mechanistic and therapeutic study, including the roles these factors may play in priming the TME for invasion. It is also important to elucidate specific TAM subsets or polarization states beyond the traditional M1/M2 model responsible for the secretion of pro-invasive factors, as well as identifying the TME features driving the presence of these key TAM polarization states and subtypes. Regional variations in TAM-GBM interactions will also be important to study given the microenvironmental differences between the tumor core and edge. Finally, understanding how TAM infiltration and function relates to GBM subtype and tumor recurrence would focus efforts to target TAMs in patients who would most benefit.

## Methods

### Me-HA synthesis

HA hydrogels were synthesized as previously described.^65^ Briefly, methacrylic anhydride (Sigma-Aldrich, 94%) was used to functionalize sodium hyaluronate (Lifecore Biomedical, Research Grade, 66 kDa – 99 kDa) with methacrylate groups (Me-HA). The extent of methacrylation per disaccharide was quantified by ^1^H nuclear magnetic resonance spectroscopy (NMR) and found to be ∼85% for materials used in this study. To add integrin-adhesive functionality, Me-HA was conjugated via Michael Addition with the cysteine-containing RGD peptide Ac-GCGYGRGDSPG-NH2 (Anaspec) at a concentration of 0.5Lmmol/l.

### HA hydrogel crosslinking

To form hydrogels, 6 wt. % Me-HA conjugated with 0.5 mM integrin binding peptide (RGD) was crosslinked with a protease-cleavable peptide (KKCG-GPQGIWGQ-GCKK, Genscript) in phenol red-free serum-free Dulbecco’s Modified Eagle’s Medium (DMEM, Thermo Fisher Sci, 21-063-029) containing GBM cell spheroids. Unless otherwise stated, all experiments in this study utilized a final 1.5 wt.% Me-HA hydrogel crosslinked with 3.064 mM peptide to yield a shear modulus ∼200 Pa, which is within the range of values typically reported for brain tissue.^66–68^

### Hydrogel rheological characterization

Hydrogel stiffness was characterized by shear rheology via a Physica MCR 301 rheometer (Anton Paar) with an 8-mm parallel plate geometry for γ = 0.5% and f = 1 Hz. Frequency was controlled to be between 50 and 1 Hz for the frequency sweep at a constant strain (γ = 0.5%), and the modulus saturation curve with time was obtained under oscillation with constant strain (γ = 0.5%) and frequency (f = 1 Hz). The temperature of the gel solution was controlled (T = 37°C) with a Peltier element (Anton Paar) and the sample remained humidified throughout the experiment.

### Cell culture

GBM-43 and GBM-6 (Mayo Clinic) were cultured in DMEM (Thermo Fisher Scientific) supplemented with 10% (vol/vol) fetal bovine serum (Corning, MT 35-010-CV), 1% (vol/vol) penicillin-streptomycin (Thermo Fisher Scientific) and 1% (vol/vol) Glutamax (Thermo Fisher Scientific, 35-050-061). Bulk GBM cell lines (GBM-43, GBM-6) were harvested using 0.25% Trypsin-EDTA (Thermo Fisher Scientific) and passaged less than 30 times.

GSC-11, GSC-268, GSC-28, GSC-262, GSC-20 and GSC-295 cells were kindly provided by University of Texas M.D. Anderson Department of Neurosurgery. GSC lines were propagated as neurospheres in DMEM/F12 50/50 1X (Corning, 10-090-CV) supplemented with 2% (vol/vol) B-27 supplement (Gibco), 20 ng/mL EGF (R&D systems, 236-EG), and 20 ng/mL FGF (R&D Systems, 233-FB). All GSC cells were harvested using Accutase cell detachment solution (Innovative Cell Technologies) and passaged less than 20 times.

| Cell line | Classification | Source | References |
| --- | --- | --- | --- |
| GSC-268 | Classical | MD Anderson | [<sup>36</sup>] |
| GSC-28 | Mesenchymal | MD Anderson | [<sup>36</sup>] |
| GSC-20 | Mesenchymal | MD Anderson | [<sup>36</sup>] |
| GSC-11 | Proneural | MD Anderson | [<sup>36</sup>] |
| GSC-262 | Proneural | MD Anderson | [<sup>36</sup>] |
| GSC-295 | Proneural | MD Anderson | [<sup>36</sup>] |
| GBM-43 | Proneural | Mayo Clinic | [<sup>69</sup>] |
| GBM-6 | Classical | Mayo Clinic | [<sup>69</sup>] |

THP-1 cells (Sigma-Aldrich, 88081201-1VL) were cultured in RPMI-1640 Medium (Sigma-Aldrich, R8758-500ML) supplemented with 10% (vol/vol) fetal bovine serum (Corning, MT 35-010-CV), 1% (vol/vol) penicillin-streptomycin (Thermo Fisher Scientific), 1% (vol/vol) MEM non-essential amino acids (Thermo Fisher Scientific), 1% (vol/vol) sodium pyruvate (Thermo Fisher Scientific), 1% (vol/vol) Glutamax (Thermo Fisher Scientific, 35-050-061), 1% (vol/vol) HEPES, and 0.05mM 2-Mercaptoethanol (Sigma-Aldrich, M3148-25ML). THP-1 cells were cultured at 3 x 10^5^ cells/ml, split when reaching 8 x 10^5^ cells/ml, and passaged less than 30 times.

THP-1 cells were differentiated and polarized following previously reported protocols.^70^ In Brief, cells were differentiated at a concentration of 5 x 10^5^ cells/mL with 100 nM phorbol 12-myristate 13-acetate (PMA, PeproTech, 1652981) in complete RPMI media for 24 hours followed by 3 days of resting in complete RPMI media. Macrophages were polarized to M1 with 50 ng/ml interferon gamma (hIFN-y, Bio Basic, RC217-17) and 50 ng/ml Lipopolysacharides (LPS, Sigma-Aldrich, L2630-10MG) for 48 hours.

Macrophages were polarized to M2 with 50 ng/ml interleukin-4 (IL-4, Bio Basic, RC212-15-5) for 48 hours (**Ext Fig 1B**). M1/M2 polarization was confirmed by measuring mRNA expression levels of established M1 and M2 markers with qPCR (**Ext Fig 1C,D)** as well as by noting morphological changes associated with differentiation and polarization **(Ext Fig 1E)**.

HMC3 cells (ATCC, CRL-3304) were cultured in DMEM/F12 50/50 1X (Corning, 10-090-CV) supplemented with 10% (vol/vol) fetal bovine serum (Corning, MT 35-010-CV) and 1% (vol/vol) penicillin-streptomycin (Thermo Fisher Scientific). Microglia were polarized to M1 with 50 ng/ml interferon gamma (hIFN-y, Bio Basic, RC217-17) and 50 ng/ml Lipopolysacharides (LPS, Sigma-Aldrich, L2630-10MG) for 48 hours. Microglia were polarized to M2 with 50 ng/ml interleukin-4 (IL-4, Bio Basic, RC212-15-5) for 48 hours.

Cells were seeded at a density of 3x 10^4^ cells/cm^2^, split when confluency reached 90%, and passaged less than 20 times. All cells were screened every six months for mycoplasma and validated every year by Short Tandem Repeat (STR) analysis at the University of California Cell Culture Facility. For experiments in hydrogels, 0.5% penicillin and streptomycin (Gibco, 15140122) and 0.125 µg/ml Amphotericin B solution (Sigma-Aldrich, A2942) was added to cell culture media.

### Conditioned media collection

To prepare Conditioned Media (CM) cells were cultured in DMEM/F12 50/50 1X (Corning, 10-090-CV) at a density of 1M cells/ml. After 48 hours, CM collected and centrifuged at 130 *g* for 5 minutes to remove dead cells and debris and passed through a 25mm 0.45µm sterile cellulose acetate filter (VWR - 76479-040). CM was used immediately or stored in −20°C prior to use. For indirect co-culture experiments, cell culture supplements (B-27 or FBS) were added to CM immediately before use. Multiple freeze/thaw cycles were avoided.

### Heat treatment of CM

CM was harvested as described above. Samples were placed in a dry heat bath set at 100°C. Samples were incubated for 1 hour and cooled to room temperature before use. If applicable, cell culture supplements (B-27 or FBS) were added to CM immediately before use.

### EV purification

We used ultracentrifugation and Nanoparticle Tracking Analysis to isolate and characterize M2-polarized macrophage-derived EVs.Conditioned media (CM) was harvested as described above. Microvesicles were removed by centrifugation at 5,000xg for 15 minutes. Supernatant was collected and ultracentrifuged at 140,000xg (28,000 RPM) for 1.5 hours in a SW-28 rotor (Beckman Coulter). The supernatant was collected (soluble protein fraction) and the crude EV pellet was resuspended in 1X PBS and ultracentrifuged at 160,000xg for 2 hours. The supernatant was discarded, and the crude EV pellet was resuspended in 0.02 µm filtered PBS for further characterization.

### Nanoparticle tracking analysis

EV sizes and quantities were estimated using the NanoSight NS300 instrument equipped with a 405-nm laser (Malvern Instruments), analyzed in the scatter mode. Silica 100-nm microspheres (Polysciences) served as a control to check instrument performance. Vesicles collected as described above were diluted 1,000× with 0.02 µm filtered PBS. The samples were introduced into the chamber automatically, at a constant flow rate of 50 during five repeats of 60-s captures at camera level 13 in scatter mode with Nanosight NTA 3.1 software (Malvern Instruments). The size was estimated at detection threshold 5 using Nanosight NTA 3.1, after which “experiment summary” and “particle data” were exported. Particle numbers in each size category were calculated from the particle data, in which “true” particles with track length >3 were pooled, binned, and counted with Excel (Microsoft). M2 CM contained EVs at a concentration of 1.67 x 10^8^ ± 7.47 x 10^6^ particles/mL with 100-400 nm diameters (mean: 229 ± 89.3 nm, mode: 178.5 nm) according to Nanoparticle Tracking Analysis of concentrated EVs (**Fig. 3c**). For the invasion assay, EVs were reconstituted in GSC maintenance media at a concentration of 1.67 x 10^8^ particles/mL, the mean concentration of EVs in M2 CM.

### Generation of spheroids

Tumorspheres were fabricated using Aggrewell Microwell Plates (Stemcell Technologies). Briefly, 1.2E5 cells were seeded into a single well of the Aggrewell plate to form spheroids consisting of 100 cells after 48 hour incubation at 37°C and 5% CO_2_.

### 3D spheroid invasion assay

Spheroids were resuspended in phenol red-free serum-free Dulbecco’s Modified Eagle’s Medium (DMEM, Thermo Fisher Sci, 21-063-029) at a density of 1.5 spheroids/ul and used as solvent for HA hydrogel crosslinking. For direct co-culture experiments, THP-1-derived macrophages were added to DMEM solvent for a final concentration of 3.5k cells/µl hydrogel. For invasion analysis in tumorsphere invasion assays, spheroids were imaged every 2 days using Eclipse TE2000 Nikon Microscope with a Plan Fluor Ph1 ×10 objective. Images were acquired using NIS-Elements Software. For each spheroid, total spheroid area was outlined in ImageJ and normalized to total day 0 spheroid area. For experiments with direct co-culture of GBM and THP-1 cells, GSCs were transduced with CAG-GFP lentivirus (Cellomics, PLV-10057-50) and selected with 1 µg/ml puromycin (Invitrogen, A1113803) prior to experiments.

### Invasion device fabrication

Devices were fabricated following a modified version of our previously published invasion device protocol.^28^ In brief, an acrylic mold was made by laser-cutting and stacking 1.5 mm-thick acrylic posts and acrylic slide (CLAREX Precision Thin Sheet, 1.5 mm, Astra Products) on top of a glass microscope slide (Fisherbrand, 12-550-A3). Next, 0.00695 mm outer diameter cleaning wires (Hamilton, 18302) were inserted into the double channel posts to serve as wire and syringe guides during hydrogel casting and cell seeding. Polydimethylsiloxane (PDMS) was fabricated by mixing a 10:1 mass to mass ratio of Sylgard 184 elastomer with the initiator (Dow Corning) and the mixture was pipetted into the acrylic mold and wire assembly and cured at 80°C for 2 hours. After curing, the wires, acrylic slide, and acrylic posts were carefully removed. A razor blade with an acrylic guide was used to cut a 5 mm by 75 mm rectangle in the center of the PDMS and the acrylic laser cut lid was attached to the PDMS and glass slide with epoxy. New 0.00695 mm outer diameter cleaning wires were reinserted into the PDMS guides and the entire device was UV-treated for 10 min and stored in cold room prior to use. Laser cutter designs are available upon request.

On the day of experiment, devices were brought to room temperature and the HA hydrogel solution was casted within the acrylic molds around the wire and incubated for 1 hour in a humidified 37°C chamber. After crosslinking, devices with hydrogels and wire were submerged in cell culture media for at least 10 min, before removal of the wire which left an open channel. Afterwards, 100k single cells were seeded into each open channel and the wires were reinserted into the device to plug each end of the cell reservoir. The entire device was placed into a 4-well plate and bathed in 10 ml of medium. Medium was changed every 3 days.

For quantification of invasion in devices, cells in devices were imaged every 7 days using Eclipse TE2000 Nikon Microscope with a Plan Fluor Ph1 ×10 objective. Images were acquired and stitched using NIS-Elements Software. For each device, total cell area was outlined in ImageJ and normalized to total day 0 cell area.

### Device Assembly and RNA extraction

For RNA extraction of cells on tissue culture plastic, phenol-free total RNA was extracted from cells directly in cell culture wells using RNeasy Plus Micro Kit with gDNA eliminator columns (Qiagen) following the manufacturer’s protocol. For RNA extraction of cells in invasion devices, devices were carefully disassembled, and if applicable, the invasive cells were physically separated from the non-invasive “core” cells based on distance from channel using a scalpel. The invasive cell populations isolated from devices cultured in GSC maintenance media were not large enough to submit for bulk RNA-seq independently and were pooled with the core cell fractions from control devices. For RNA extraction of tumorsphere invasion assays, hydrogels and cells were placed in Eppendorf tubes. All samples were treated with 10k U/ml Hyaluronidase from bovine testes, Type IV-S (Sigma-Aldrich, H3884) for 15 min with agitation until the hydrogel was fully degraded. Afterwards, RNA was extracted using TRIzol Reagent (Invitrogen, 15596018) according to manufacturer’s recommendations. In brief, 1M cells were mixed with 100 µl TRIzol Reagent and the solution was vortexed for 30 seconds. 20 µL chloroform (Sigma, c2432) was added to the mixture and the solution was placed on ice for 15 minutes. Sample were centrifuged at 14000 rpm for 10 minutes and the clear RNA layer was carefully transferred to a new Eppendorf tube. Each RNA sample was mixed with a 1:1 volume of 2-Propanol (Sigma-Aldrich, 190764-4L) and 0.5 µL Glycogen, RNA grade (Thermo Scientific, R0551). After an overnight incubation at −20°C, samples were centrifuged at 14000 rpm for 10 minutes at 4°C and the RNA pellet was rinsed twice by aspirating the supernatant, adding 400 µL 75% Ethanol and centrifuging at 14000 rpm for 1 minute. Following the second rinse, the supernatant was carefully and completely removed, and RNA pellets were incubated at room temperature until visibly dry. RNA was dissolved in 30 µL RNase-free DNase-free distilled H_2_O and RNA concentration and purity was measured with Nanodrop.

### cDNA synthesis

cDNA was synthesized from extracted RNA using iScript cDNA Synthesis Kit (BioRad, #1708891) following the manufacturer’s protocol.

### RNA-sequencing and differential gene expression analysis

Isolated RNA was sent to Novogene Corporation Inc. (Sacramento, CA) for library construction, quality control and sequencing. In brief, messenger RNA was purified from total RNA using poly-T oligo-attached magnetic beads. After fragmentation, the first strand cDNA was synthesized using random hexamer primers followed by the second strand cDNA synthesis. The library was ready after end repair, A-tailing, adapter ligation, size selection, amplification, and purification. The library was checked with Qubit and real-time PCR for quantification and bioanalyzer for size distribution detection. Quantified libraries will be pooled and sequenced on Illumina platforms, according to effective library concentration and data amount. Data was filtered to remove low-quality reads and sequences containing adapters and differential expressed genes were identified using DESeq2. Statistically significant differentially expressed genes were chosen based on a statistical cutoff of |Log2FC| ≥ 1 and adjusted p-value < 0.05.

### Pathway enrichment analysis

We used Enrichr^71–73^ to perform enrichment analysis of gene lists using the Molecular Signatures Database (MSigDB^74^) hallmark gene set collection. MSigDB Pathways were ranked by Odds Ratio.

### Receptor-ligand interaction analysis

To infer receptor-ligand interactions between GSCs and M1/M2 THP-1 macrophages, we compared our RNA-seq data of GSC-20s and our global proteomics data of M1/M2 macrophage CM to a database of 3631 receptor-ligand interactions ^75,76^ which was downloaded and analyzed in RStudio (RStudio, Inc., 2022.12.0+353). We first filtered the M1/M2 macrophage CM dataset to identify conventionally secreted proteins using The Human Protein Atlas (<u>Predicted: secreted</u>) database of predicted secreted proteins.^39^ We then annotated cognate pairs co-expressed by GSC-20s for which the Base Mean transcript expression of the receptor in the GSC-20 + M2 CM (invasive fraction) sample is ≥1. This produced 255 receptor-ligand pairs for which the receptor is expressed by GSC-20s and the ligand is expressed by M1/M2 macrophages.

To identify the receptor-ligand interactions dependent on macrophage polarization state, we grouped our inferred receptor-ligand pairs into M2- and M1-specific interactions using Fold Change (FC)=(M2 normalized intensity)/(M1 normalized intensity). FC ≥ 2 indicated M2-specific interactions, 2<FC>0.5 indicated pan-macrophage interactions, and FC≤0.5 indicated M1-specific interactions. This analysis yielded 100 M1-specific interactions, 97 M2-specific interactions, and 58 pan-macrophage interactions (**Dataset 3**).

### Acquisition of scRNA-seq datasets

Differential expression genes from scRNA-seq analysis of 9 patient GBMs was downloaded from <u>PMID: 34434898</u> (ref. ^77^) which obtained 10 X genomics scRNA-seq data from the Gene Expression Omnibus (GEO) (https://www.ncbi.nlm.nih.gov/geo/) database under <u>GSE131928</u>.^40^ The sequencing analyses were conducted on samples from nine fresh GBM tumors from adult patients in Massachusetts General Hospital (MGH). Dimensionality reduction, unsupervised clustering, and identification of cell types and marker genes in different clusters were performed as previously described.^77^ In brief, single-cell transcriptomes for a total of 16,201 cells were retained after initial quality controls and were differentiated into 22 different clusters, which were further sorted into three main cell types by comparing their gene expression characteristics using the following marker genes: malignant cells (*GAP43, GPM6B, SEC61G, PTN*), tumor-associated macrophages (*C1QA, C1QB, C1QC, TYROBP, CD68*) and T lymphocytes (*CD3G, GZMH, IL2RB, PR1F, and ICOS*). Pct.1 represents the percentage of cells where the gene is detected in specific cluster and pct.2 represents the percentage of cells on average in all the other clusters where the gene is detected.

Enrichment Score was calculated by dividing pct.1 by pct.2. Clusters 2,3,6,7,9, and 19 represent tumor-associated macrophages, cluster 17 represents T-lymphocytes, and clusters 0,1,4,5,8,10-16,18,20, and 21 represent malignant cells. To comprehensively characterize the signatures of TAMs in GBM, the cells in the macrophage and microglia groups were subclustered and reanalyzed. In total, 5,896 cells were sorted and hierarchically sorted into 14 clusters, as described in ref. ^77^. Clusters 4,5,6,7,10 and 11 represent resident microglia TAMs and clusters 0,1,2,3,8,9 and 13 represent bone marrow-derived TAMs.

Differential expression genes from human blood-derived and microglial tumor-associated macrophages were downloaded from <u>PMID: 29262845</u>, which performed scRNA-seq on a cohort of 19 total patients comprising 5,455 TAMs (DESeq adj. p value < 1e-3).^41^ The scRNA-seq datasets were mined for genes encoding proteins that were detected by mass spectrometry in M2 CM and identified in our GSC-macrophage interaction analysis. Data was downloaded and analyzed in RStudio.

### Acquisition of The Cancer Genome Atlas (TCGA) gene expression

TCGA data was queried and downloaded using GlioVis^42^ data portal for visualization and analysis of brain tumor expression datasets. Data was graphed and analyzed in Prism GraphPad.

### Quantitative PCR (qPCR)

qPCR was carried out using Applied Biosystems PowerUp SYBR Green Master Mix (Thermo Scientific, A25918) and primers following the manufacturer’s recommended protocol in a BioRad CFX Connect Real-Time PCR cycler with a 5 μM final forward and reverse primer concentration. Ct values were calculated using CFX Maestro software accompanying the real-time cycler and relative gene expression was calculated using the 2^(-delta delta Ct) method. Each gene expression was internally normalized by the expression level of housekeeping gene run on the same qPCR batch. Primer sequences were designed using either Primer-BLAST^78^ or PrimerBank.^79^ Primer sequences are listed in **Dataset 4**.

### Mass spectrometry sample preparation

To prepare Conditioned Media (CM) for mass spectral analysis, M1- and M2-polarized macrophages were cultured in DMEM/F12 50/50 1X (Corning, 10-090-CV) at a density of 1M cells/ml. After 48 hours, CM from M1- and M2-polarized macrophages was collected and centrifuged at 130 *g* for 5 minutes to remove dead cells and debris. The CM was then added to Amicon Ultra-4 Centrifugal Filter Unit tubes (Millipore, UFC8010-24) and concentrated by centrifugation at 10,000 *g* for 15 minutes resulting in a 20-fold reduction in volume (V_i_= 4.0 mL; V_f_ = 0.2 mL).

In preparation for mass spectral analysis, samples underwent a trypsin in-solution protein digest followed by sample desalting. Peptides were analyzed in biological triplicate on a ThermoFisher Orbitrap Fusion Lumos Tribid mass spectrometry system equipped with an Easy nLC 1200 ultrahigh-pressure liquid chromatography system interfaced via a Nanospray Flex nanoelectrospray source.

Samples were injected on a C18 reverse phase column (25 cm x 75 µm packed with ReprosilPur C18 AQ 1.9 µm particles). Peptides were separated by an organic gradient from 5 to 30% ACN in 0.02% heptafluorobutyric acid over 180 min at a flow rate of 300 nl/min. Spectra were continuously acquired in a data-dependent manner throughout the gradient, acquiring a full scan in the Orbitrap (at 120,000 resolution with an AGC target of 400,000 and a maximum injection time of 50 ms) followed by as many MS/MS scans as could be acquired on the most abundant ions in 3 s in the dual linear ion trap (rapid scan type with an intensity threshold of 5000, HCD collision energy of 32%, AGC target of 10,000, maximum injection time of 30 ms, and isolation width of 0.7 *m/z*). Singly and unassigned charge states were rejected. Dynamic exclusion was enabled with a repeat count of 2, an exclusion duration of 20 s, and an exclusion mass width of ±10 ppm.

### Mass Spectrometry analysis

Protein identification and quantification were done with Integrated Proteomics Pipeline (IP2, Integrated Proteomics Applications, Inc. San Diego, CA) using ProLuCID/Sequest, DTASelect2 and Census.^80–83^ Identified proteins that were detected in only one replicate per sample were filtered out from dataset. Protein fold change was calculated by dividing the average normalized M2 intensity by the average normalized M1 intensity for each protein.

### Additional reagents

| **Cell Culture Reagents** |  |  |  |
| --- | --- | --- | --- |
| Reagent | Product | Vehicle | Concentration |
| rhBIGH3 | R&D Systems **3409-BG** | 1X PBS | 20 ng/ml |
| rhS100A9 | R&D Systems, **9254-S9** | dH2O | 5 μg/ml |
| rhTGFβ1 | R&D Systems, **240-B** | 4 mM HCl + 1 mg/mL BSA | 20 ng/ml |
| Temsirolimus | MedChemExpress, HY-50910 | DMSO | 10 μM |
| MHY1845 | SelleckChem, S7811 | DMSO | 10 μM |
| ABR-238907 | MedChemExpress, HY-141537 | DMSO | 25 μM |
| Reagent | Product | Host | Concentration |
| anti-BIGH3 | Proteintech, **10188-1-AP** | Rabbit | 10 μg/ml |
| anti-S100A9 | Abcam, ab63818 | Rabbit | 10 μg/ml |
| IgG | Sigma-Aldrich, I5006 | Rabbit | 10 μg/ml |

| **Immunocytochemistry Reagents** |  |  |
| --- | --- | --- |
| Reagent | Product | Concentration |
| DAPI | Roche, 10236276001 | 1 μg/ml |
| Anti-KI-67 [SP6] | Abcam, ab16667 | 0.15 μg/ml |

### Statistical analyses

Unless otherwise specified, statistical analyses were performed on Prism GraphPad v.9.0.2. Statistical tests used two-tailed one-way ANOVA and post-hoc tests for multiple comparisons, or Student’s t-test or Wilcoxon test for comparison between two groups. Box and whiskers represent minimum to maximum. Error bars represent standard deviation. For all statistical tests unless otherwise noted: \**P* < 0.05, \*\**P* < 0.01, \*\*\**P* < 0.001, \*\*\*\**P* < 0.0001.

## Acknowledgements

This work used the Vincent J. Coates Proteomics/Mass Spectrometry Laboratory at UC Berkeley, supported in part by NIH S10 Instrumentation Grant S10RR025622. Glioma Stem Cell lines were developed at the University of Texas M. D. Anderson Department of Neurosurgery support by grants from the National Cancer Institute (1R01CA214749, 1R01CA247970, P30CA016672 and 2P50CA127001) and the University of Texas M. D. Anderson Moon Shots Program^TM^. We also thank Dr. Randy Schekman (UC Berkeley) for providing the NanoSight NS300 instrument, Liang Ma for advising on Extracellular Vesicle isolation and Kwasi Amofa for providing qPCR primers.

This work was supported by awards from the National Science Foundation (Graduate Research Fellowship to E.A.A.) and the National Institutes of Health (R01CA227136 and R01CA260443 to M.K.A. and S.K.).

## Author Contributions

E.A.A, S.K. and M.K.A. conceived the project. E.A.A. wrote the manuscript, performed most of the experiments and prepared the figures. D.W. contributed to the experimental work. All authors contributed to editing and revising the manuscript.

## Supplementary Data

**Dataset 1.** Bulk RNA-seq differentially expressed genes

**Dataset 2.** Proteins identified by mass spectrometry in conditioned media isolated from M1- and M2-polarized THP-1 macrophages

**Dataset 3.** Receptor-ligand pairs identified from Receptor-ligand analysis of GSC-20 transcriptome and THP-1 macrophage proteomics

**Dataset 4.** qPCR primer sequences

## EXTENDED FIGURES

**Ext. Fig. 1:**
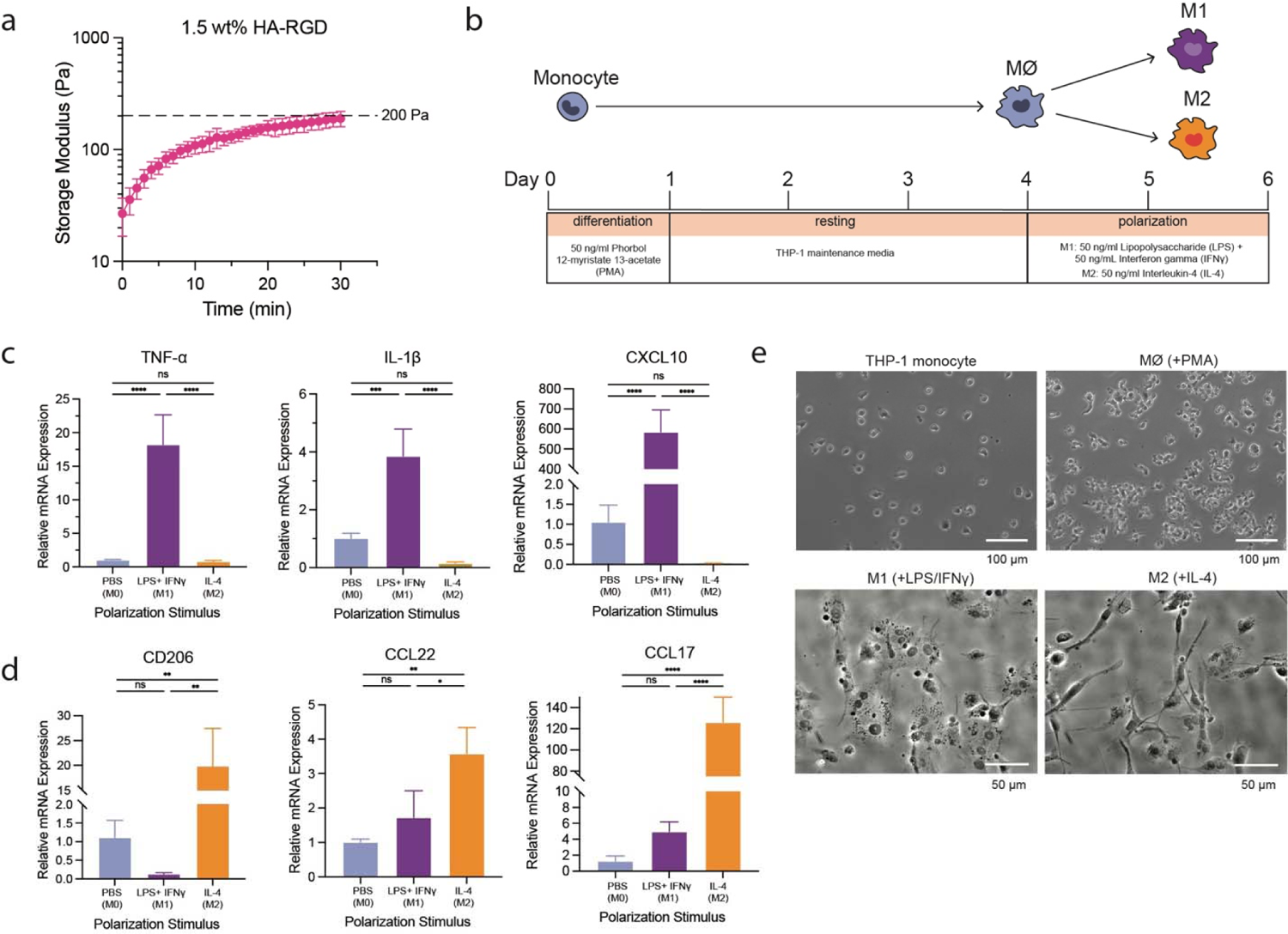
Hydrogel characterization and THP-1 macrophage differentiation and polarization validation. **a**, Hydrogel storage modulus (G’) (in Pascals) versus time after the crosslinker is added. Time=0 represents the intersection of storage modulus (G’) and loss modulus (G’’) (n=3 hydrogels). **b,** Schematic of THP-1 differentiation and polarization timeline. **c,d,** qPCR gene expression of (**c**) M1 and (**d**) M2 markers in THP-1-derived macrophages polarized with LPS + IFNγ or IL-4. **e,** Representative phase images of THP-1 monocytes/macrophages throughout differentiation and polarization process.

**Ext. Fig. 2:**
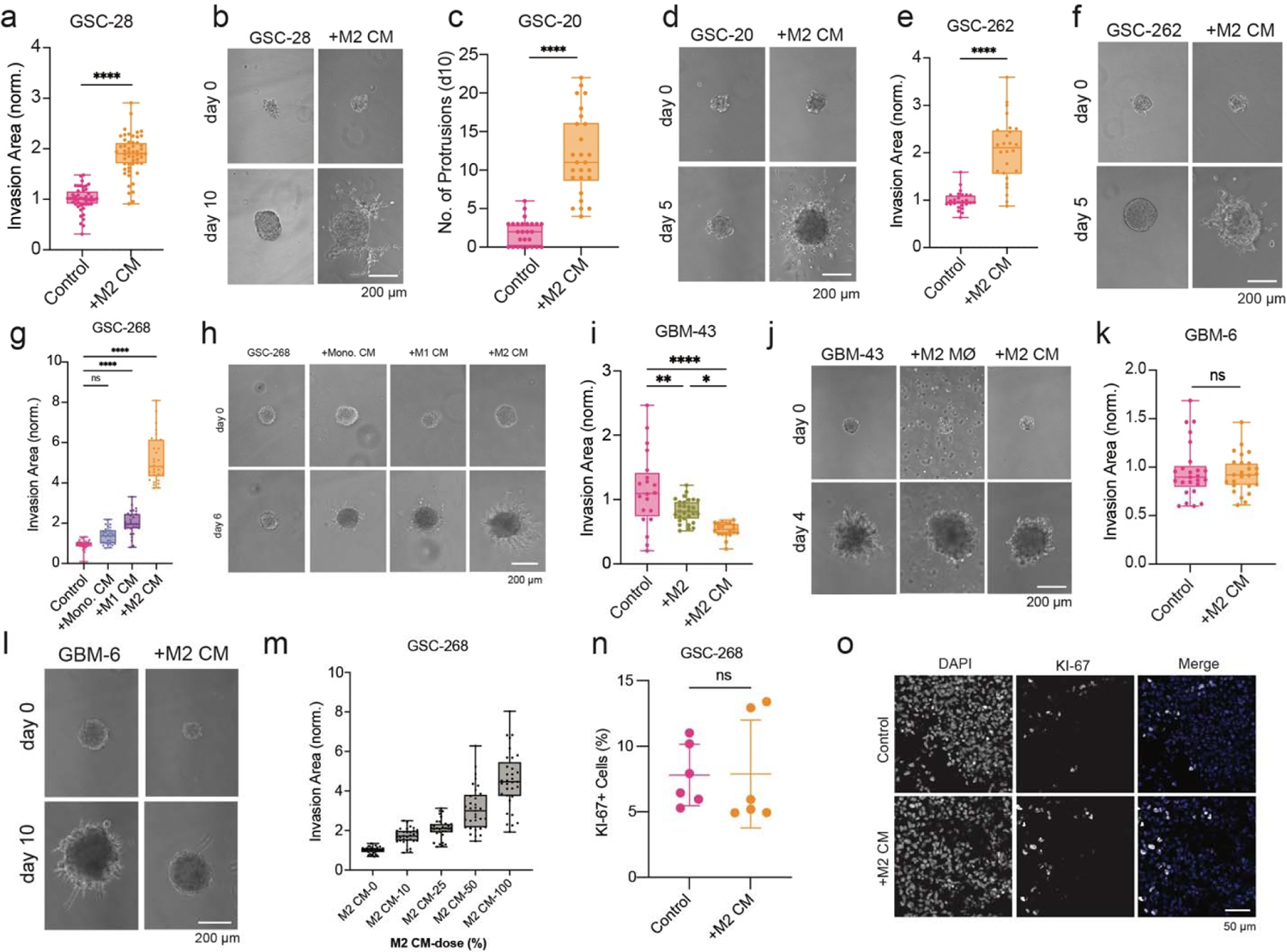
Validation of M2 macrophage-induced invasion across multiple cell lines. **a,b**, GSC-28 invasion assay with M2 CM (**a**) quantification and (**b**) representative phase images. **c,d,** GSC-20 invasion assay with M2 CM (**c**) quantification and (**d**) representative phase images. **e,f,** GSC-262 invasion assay with M2 CM (**e**) quantification and (**f**) representative phase images. **g,h,** GSC-268 invasion assay with CM from monocytes (mono.), M1 macrophages, and M2 macrophages (**g**) quantification and (**h**) representative phase images. **i,j,** GBM-43 invasion assay with direct M2 macrophage co-culture and M2 macrophage conditioned media (CM) (**i**) quantification and (**j**) representative phase images. **k,l,** GBM-6 invasion assay with M2 CM (**k**) quantification and (**l**) representative phase images. **m,** GSC-268 invasion assay with varied doses of M2 CM quantification through serial dilutions of M2 CM. **n,o,** KI-67 staining of fixed GSC-268 cells in 3D invasion assay (**n**) quantification and (**o**) representative immunofluorescent images.

**Ext. Fig. 3:**
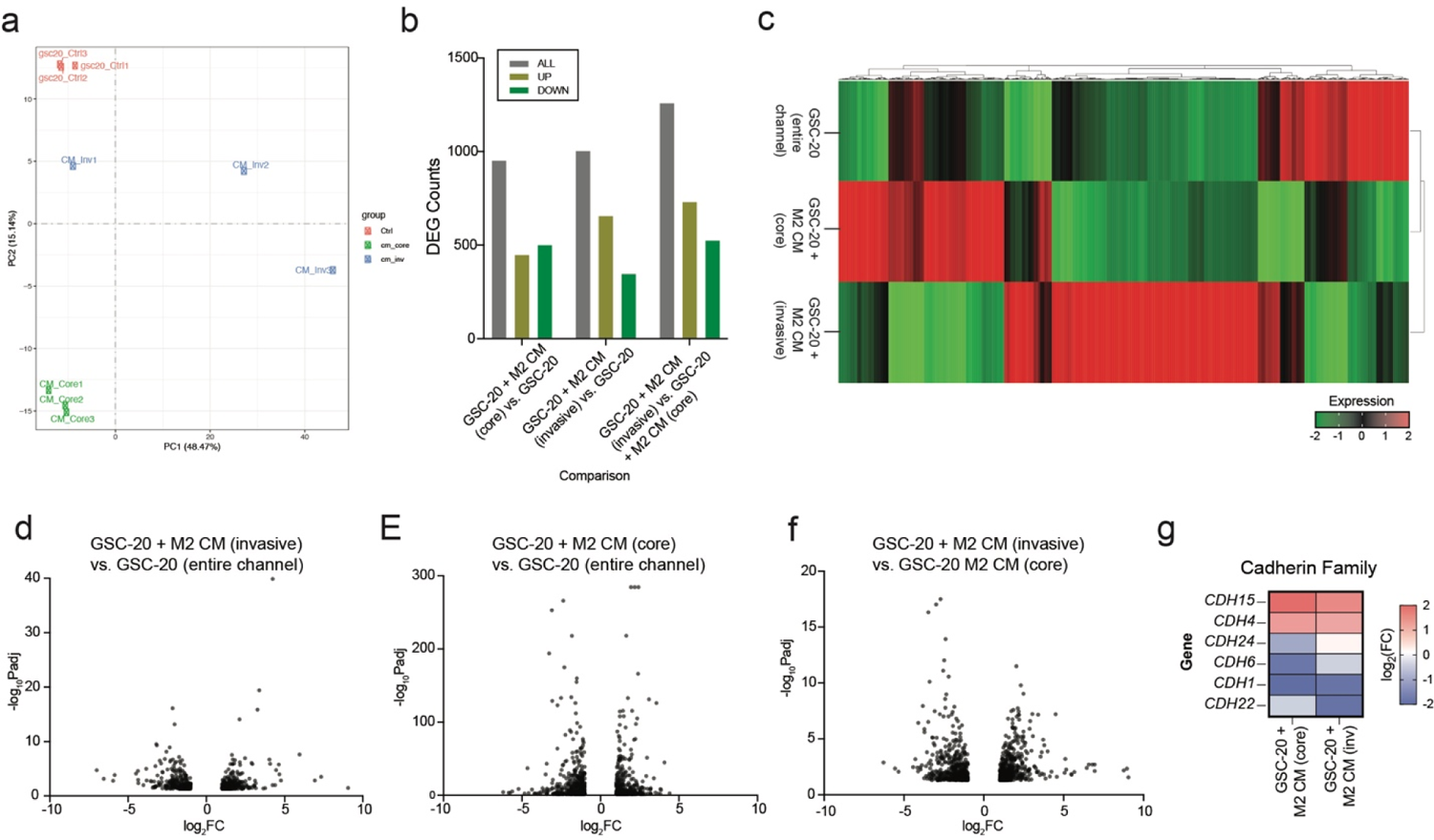
Bulk RNA-seq quality control and additional analysis. **a**, Principal Component Analysis (PCA) of bulk RNA-seq samples. **b,** Bar graph showing up, down, and total differentially expressed gene (DEG) counts for each pairwise comparison using cut offs abs(log2FC) > 1 and *P*_adj_ < 0.05. **c,** Heat map illustrating relative differential gene expression changes across samples. **d-f,** Volcano plots of DEGs for (**d**) GSC-20 invasive fraction of M2 CM devices and entire channel of GSC-20 control devices, (**e**) GSC-20 core fraction of M2 CM devices and entire channel of GSC-20 control devices and (**f**) GSC-20 invasive fraction of M2 CM devices and GSC-20 core fraction of M2 CM devices. **g,** Heat map showing GSC-20 relative gene expression levels of cadherin family genes obtained by bulk RNA-seq of cells isolated from invasion devices. Expression levels normalized to GSC-20 devices in control media (entire channel).

**Ext. Fig. 4:**
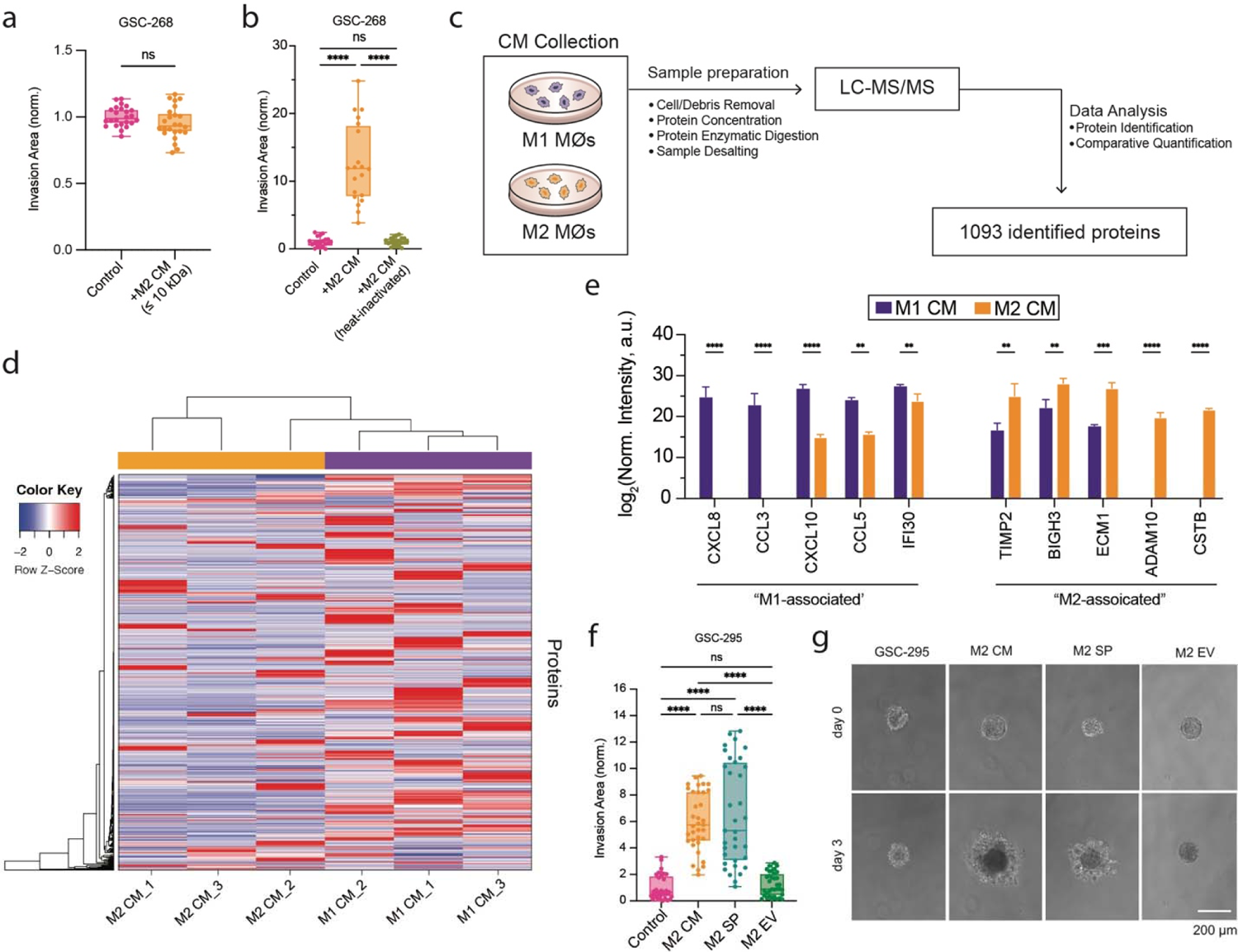
Mass Spectrometry quality control and additional analysis. **a**, GSC-268 invasion assay quantification using size-based filtered M2 CM (≤ 10 kDa fraction). **b,** GSC-268 invasion assay quantification using heat-inactivated M2 CM. **c,** Schematic of mass spectrometry sample collection, preparation, and analysis. **d,** Heat Map illustrating differential protein expression across samples. **e,** Relative protein intensity of M1- and M2-associated proteins identified in M1 and M2 CM. **f,g** GSC-295 invasion assay with M2 CM, M2 soluble proteins (SP) and M2 extracellular vesicles (EV) (**f**) quantification and (**g**) representative phase images.

**Ext. Fig 5:**
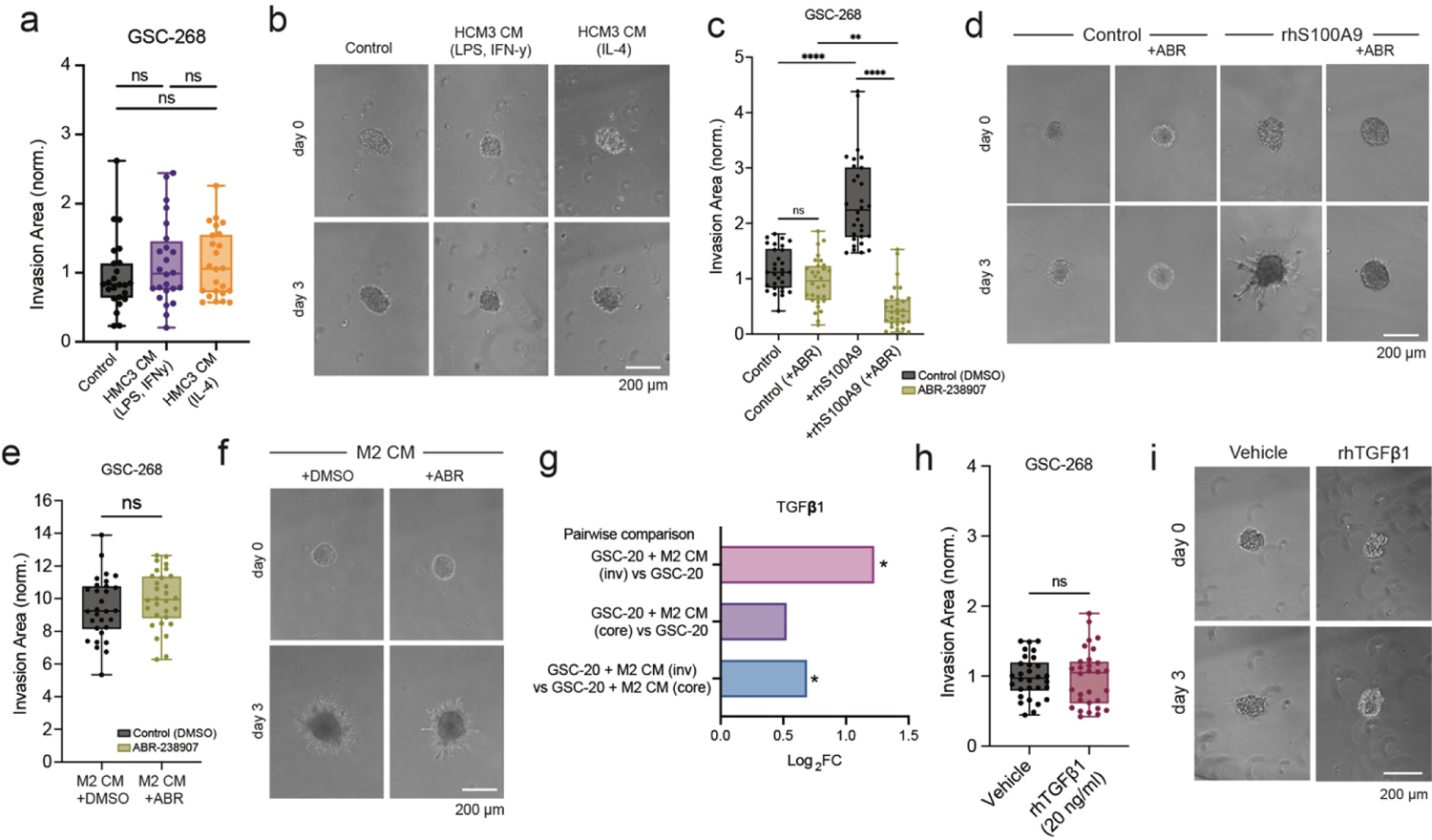
Validation of TAM-derived BIGH3 specificity and intracellular pathway. **a,b**, GSC-268 invasion assay with HMC3 CM (labeled with polarization stimuli) (**a**) quantification and (**b**) representative phase images. **c,d,** GSC-268 invasion assay with 5 µg/ml rhS100A9 and 25 µM ABR-238907 (ABR) (**c**) quantification and (**d**) representative phase images. **e,f,** GSC-268 invasion assay with M2 CM and 25 µM ABR-238907 (ABR) (**e**) quantification and (**f**) representative phase images. **g,** Bar plot showing differential TGFβ1 gene expression across sample comparisons from bulk RNA-seq dataset. **h,i,** GSC-268 invasion assay with 20 ng/ml rhTGFβ1 (**e**) quantification and (**f**) representative phase images.

